# Discordant expression profile between RNA and protein for the genes involved in immune response network in adenovirus type 2 infected cells

**DOI:** 10.1101/302851

**Authors:** Hongxing Zhao, Maoshan Chen, Alberto Valdés, Sara Bergström Lind, Ulf Pettersson

## Abstract

Alternation of cellular genes expressions during Adenovirus type 2 (Ad2) infection in IMR-90 cells was studied using paired-end sequencing and stable isotope labeling of amino acids in cell culture mass spectrometric analysis (SILAC-MS). At transcriptional level, cellular genes involved in different pathways revealed distinct expression profiles. At early phase, the genes involved in regulation of cellular immune response, cellular signaling and cell growth control were among the most deregulated. Later follows, in an orderly fashion, genes involved in cell cycle control, DNA replication and further on genes engaged in RNA processing and protein translation. Comparison of cellular gene expression at transcriptional and posttranscriptional levels revealed low correlation. Here we highlight the genes which expose opposite expression profiles with an emphasis on key factors that play important roles in cellular immune pathways including NFκB, JAK/STAT, caspases and MAVS. Transcription of many of these genes was transiently induced early, but became down-regulated in the late phase. In contrast, their expressions at protein level were up-regulated early and so sustained until late phase of infection. Suppression at the transcriptional level and enhancement at the protein level of immune response genes most likely illustrate counteractions between Ad2 and its host cell.

**Importance:** Our paper comprises a state of the art quality transcriptomics data set unravelling the alterations in gene expression that take place during different phases of an adenovirus infection. The information allows us to draw conclusion about the cellular pathways that are perturbed by the virus. The data set also provides an important resource for scientists in general for future studies on mechanisms behind host/virus interactions in efforts to design tools for combatting virus infections.

Moreover, our paper includes novel proteomics information unravelling an unexpected role of post transcriptional events in cellular gene expression, demonstrating that the current picture of the adenovirus replication cycle is simplified.

## Introduction

Human adenovirus (Ad) infection leads to alternations of host cell gene expression and biosynthetic processes. It is a stepwise, but efficient mode of turning host transcriptional, translation and metabolism to facilitate the replication of adenovirus. Most interactions between host cell and virus take place during the early phase. Adenovirus-mediated regulation of cellular gene expression emphasizes two major aspects: interference with host defense mechanisms and induction of its host cell to enter S-phase of the cell cycle. It has also been shown that cells are reprogrammed epigenetically as a result of adenovirus early-region function at different times after infection (1). Adenovirus expresses several regulatory proteins from early regions 1A (E1A), E1B, E3, and E4. E1A is the first viral gene expressed and plays essential roles in regulation of viral and cellular genes expression (2). E1A proteins are crucial for the induction of the S phase of the cell cycle, cell proliferation and cell transformation through its ability to target different cellular transcriptional regulators, such as pRb, p300/CBP, CtBP, p400/TRRAP (3–10). E1A proteins also interfere with host immune response by blocking type I IFN-inducible gene expression (11), as well as by preventing the peptide presentation to the immunoproteosome by interacting with MECL1 (12). E1B encodes two major proteins, the E1B-55K and E1B-19K proteins. E1B-55K is a multi-functional protein and plays a major role in counteracting the cellular proapoptotic program. Association of E1B-55k and E4 orf6 proteins with several cellular proteins, Cullin 5, TCEBs and RBX1 forms a virus-specific E3 ubiquitin ligase which then targets specific cellular proteins for degradation (13). The E1B-55K protein serves as the substrate-recognition subunit via distinct sequences and targets the p53 protein, thereby promoting degradation of p53 (14, 15). The E1B-19K protein is a viral Bcl-2 homologue that acts as a broad inhibitor of mitochondria-dependent apoptosis (16, 17). It interferes directly with the activity of p53 when translocated into the mitochondria (18). Proteins generated from the E3 region also play a very important role in countering host antiviral defenses (19). E3-gp19K prevents the exposure of viral peptides on the cell surface by blocking the transport of the class I major histocompatibility complex (MHC I) molecule to the cell surface and the loading of peptides by tapasin (20–22). The E3-10.4K and 14.5K (RIDα/β) complex inhibits tumor necrosis factor alpha (TNFα) and Fas ligand-induced apoptosis through internalization and degradation of the death domain containing receptors (23). In addition, the E3-10.4K/14.5K complex blocks the activation of NFκB by preventing it from entering the nucleus and inhibiting the activity of the kinase complex IKK (24). Proteins encoded by the E4 region are involved in transcriptional regulation. E4 orf6/7 stabilizes the binding of E2F to the duplicated E2F binding sites in the E2 promoter (25, 26). E4 orf3 associates with E1B 55K in the nuclear promyelocytic leukemia protein oncogenic domains (POD) and reorganizes PODs during infection, thus likely involved in the regulation of transcription factor availability and activity (27). The E4 orf4 protein interacts with protein phosphatase 2A, leading to the inhibition of E1A-dependent transactivation of the junB promoter (28).

When adenovirus DNA replication commences, the infection cycle proceeds into the late phase and viral transcription changes from the early to the late pattern. E1A expression switches from a preference for the 289R transcriptional activator to the shorter E1A-243R protein (29). The E1A-243R protein functions mainly as a transcriptional repressor through its binding of p300/CBP (30). The L4-100 kDa protein, expressed from the major late transcription unit is necessary for efficient initiation of viral late mRNA translation (31–33). Furthermore, the E1B-55 kDa and E4 orf4 protein complex is involved in regulation of mRNA export from nucleus, resulting in a block of cellular mRNAs export and selective export of viral mRNAs (34, 35). As a consequence a dramatic down-regulation of cellular gene expression occurs late in infection (36).

Most studies of the adenovirus infection have been performed in Hela cells, in which adenovirus replication is very efficient and the infectious cycle is completed after 20-24 hours. Particularly, the early phase is very short, lasting for less than 6 hours. Thus, there is a narrow time window for a detailed examination of the changes of cellular gene expression. Furthermore, being transformed cells, Hela cells grow rapidly and are difficult to synchronize. Thus, genes involved in the control of cell cycle and growth might escape detection. Therefore, human primary cells, like human lung fibroblasts (IMR-90) or foreskin cells (HFFs) have been used for a series of studies (36–40). In these cells adenovirus DNA replication starts 24 hours post infection (hpi). Based on cellular transcription profiles, an adenovirus type 2 (Ad2) infection of IMR-90 cells can be divided into four periods (36). The first period (1-12 hpi) extends from the attachment of Ad2 to the cell surface to the beginning of adenoviral early gene expression. During this time, the cellular gene expression changes are mainly triggered by the virus entry process, including attachment of the virus to cell surface receptors, and intracellular transport of the virus along microtubules. The majority of the genes deregulated during the first phase have functions linked to inhibition of cell growth and immune response. The second period covers the time from the expression of the immediate early E1A gene to the time when Ad2 DNA replication starts (12-24 hpi). During this period, there is a linear increase in the number of differentially expressed cellular genes involved primarily in cell cycle regulation and cell proliferation. The third period ranges from the beginning of DNA replication to the time when the cytopathic effect (CPE) starts (24-36 hpi). By this time, the virus has gained control of the cellular metabolic machinery, resulting in an efficient replication of the viral genome and expression of the capsid proteins. Additional changes in cellular gene expression are modest during this phase. The final period starts when CPE is apparent (after 36 hpi). The number of down-regulated genes increases dramatically and include many genes involved in intra- and extracellular structure, leading to an efficient burst of progeny.

The transcriptomics and proteomics of host cell during human adenovirus infection have been extensively studied (39–43). Early studies of cellular genes expression in adenovirus subtype C infected quiescent fibroblasts using microarray showed that genes involved in the control of cell cycle, proliferation, and growth were most highly up-regulated, while genes implicate in immune response were most significant among down-regulated genes during the early phase of infection (36, 40). With the development of high throughput sequencing technologies, the transcriptome can be explored on a genome-wide scale at single base pair resolution. A large number of differentially expressed cellular genes have been identified, and they are classified into similar gene ontology categories as identified by microarray (42). Binding sites for E2F, ATF/CREB and AP2 are prevalent in the up-regulated genes while SRF and NFkB are most dominant among down-regulated genes at two time points (12 and 24 hpi) as described in a previous study (42). Meanwhile, several proteomics approaches have been applied. Improve shotgun/bottom-up liquid chromatography-tandem mass spectrometry (LC-MS/MS)-based protein detection and quantitative techniques such as Stable Isotope Labelling of Amino acids in Cell culture (SILAC) have greatly facilitated protein identification (44, 45). These technologies have been used in studies of protein expression in an adenovirus-infected cells. Lam et al have analyzed the nucleolar proteome in Ad5-infected Hela cells (46) while Evans et al have examined the posttranscriptional stability of cellular protein in Ad5-infected Hela cells (41). Recently, a comparative proteome analysis of wild type and E1B-55K was performed to investigate the role of Ad5 E1B-55K in targeting cellular proteins with antiviral activity for proteasomal degradation (47). Furthermore, using a combined immunoaffìnity purification and LC-MS protocol, a set of 92 E1B-associated proteins were identified in Ad5-infected HFFs and it was shown that these proteins are enriched for function in the ubiquitin-proteasome system, RNA metabolism and cell cycle (48). Previously, we have presented a comparison of the cellular transcriptome and proteome of Ad2-infected IMR-90 cell at 24 and 36 hpi (39). More than 700 proteins were identified to be differentially expressed. Surprisingly, there was a very low correlation between the RNA and protein expression profiles. Here, we present a more comprehensive study of the cellular transcription profiles at four critical stages of an adenovirus infection in normal cell using paired-end sequencing. As a step further, RNA expression profiles were compared with protein expression profiles with a focus on genes involved in the cellular immune response.

## Materials and methods

### Cell culture and virus infection

Human primary lung fibroblast IMR-90 cells (American Type Culture Collection, ATCC) were initially cultured in Eagle’s minimum essential medium (EMEM) (ATCC) supplemented with 10% fetal bovine serum (FCS), 100 U/ml penicillin and 100 μg/ml streptomycin at 37 °C and 5% CO_2_. Cells were maintained in the plates for two days before infection. By fluorescence-activated cell sorting (FACS) analysis, more than 95% of the cells were characterized in G0/G1 phase. Synchronized cells were then infected with human adenovirus type 2 at a multiplicity of infection (MOI) of 100 fluorescence-forming units (FFU) in serum-free medium. Mock-infected cells were used as a control. One hour later, the medium was replaced with complete EMEM medium supplemented with 10% FBS. Infected cells were harvested and collected at 6, 12, 24, and 36 hours post infection (hpi).

### Total RNA extraction, RNA library construction and sequencing

Total RNA from infected IMR-90 cells were extracted with TRIzol® (Invitrogen), according to the manufacturer’s instructions. The quality of total RNA was evaluated with a NanoDrop 1000 spectrophotometer and an Agilent 2100 Bioanalyzer. After treatment with Ribo-Zero™ rRNA removal reagent, total RNA was used to construct cDNA library for transcriptome sequencing following the ScriptSeq™ v2 RNA-Seq library preparation kit according to the manufacturer’s protocol (Epicentre). The cDNA libraries were sequenced on a HiSeq 2000 sequencing platform (Illumina).

### Genome alignment and gene expression profile

Data cleaning was performed by removing low quality, contaminant and adapter reads from the raw reads. TopHat2 and Cufflinks were used to align the filtered reads to human Ensembl genome (http://www.ensembl.org/index.html, GRCh38) and to profile gene expression following the protocol (49), respectively. FPKM (fragments per kilobase of exon per million fragments mapped) method was employed to normalize gene expression. To strengthen the reliability of our results, lowly expressed genes (< 10 FPKM in all libraries) were filtered out.

### Identification of differentially expressed genes in Ad2-infected cells

To identify genes deregulated in early and late phases of Ad2 infection, we performed correlation analysis between samples based on normalized gene expression values using the CORREL function provided by Excel. To identify differentially expressed genes in the cells infected by Ad2, several statistical values were used. First, a fold change of a particular gene in Ad2-infected cells was calculated following the rule: fold change (Ad2-infected/mock) =y/x, while y and x represent the normalized expression values in Ad2-infected and mock cells, respectively. A cut-off of more than 2-fold increase or decreas was used. Second, a p-value that represents the significance for differential expression was calculated based on Poison distribution (50). A cut-off for p-values (< 0.05) was used for differentially expressed genes. Last, an R package called NOISeq was used to calculate the probability of differential expression of a gene in a comparison (51). Only those genes with probability > 0.7 were kept for further analysis.

### Gene Ontology and KEGG pathway enrichment

To determine the biological processes and KEGG pathways affected by human adenovirus type 2, differentially expressed genes were analyzed by DAVID Bioinformatics Resources 6.7 (http://david.abcc.ncifcrf.gov/) (52).

### SILAC-MS experiment and protein identification

The IMR-90 cells culture and protein labelling were performed as described before (36). Briefly, after growing in cell culture medium containing with heavy or light amino acids for at least six passages, cells were mock infected or infected with Ad2 at MOI of 100 FFU per cell in serum-free medium (53). A biological replicate with swapped labeling was also performed. After harvest, cells were lysed and mock- and Ad2-infected lysates of different labeling were combined in a 1:1 protein ratio. Proteins were fractionated using SDS-PAGE. Following ingel tryptic digestion (54), peptides were extracted and analyzed using nano liquid chromatography coupled on-line to a QExactive Orbitrap Plus Mass spectrometer (ThermoFisher Scientific, Bremen,Germany). Acquired data (raw-files) were imported into MaxQuant software (version:1.4.5.7) (55), and searched against a FASTA-file containing both cellular and Ad2 proteins downloaded from UniProt 2017-02. The ratio of the chromatographic areas of heavy and light peptides matching to specific proteins was used for determining the protein expression levels.

## Results and discussion

### Host cell transcriptional profiles during the course of an adenovirus infection

Alternation of cellular transcription in adenovirus infected cells has been studied before using RNA sequencing (36). However, this study included only two time points (12 and 24 hpi) in lung fibrobalst using constrained 76 bp long sequencing reads. Therefore, a more detailed study of transcription at different phases of infection using a up-graded sequencing technique is recalled. Furthermore, the correlation between transcription and protein expression need to be adressed. To this end, we have applied paired-end sequencing technology to examine the cellular RNA expression profile at 6, 12, 24 and 36 hours post infection (hpi) of IMR-90 cells. These time points represent different stages of Ad2 infection and all of our early studies on cellular various RNA expression including micro RNA (miRNA), long non-coding (lncRNA) and protein were performed under the same condition (36–38). Thus, we could correlated the expression profile between them. About 30 million 255 bp long sequence reads per sample were generated and 53-58% of them accounted for mRNA. From them 6,860 cellular genes were identified to be transcribed to a significant level with a minimum of 10 FPKM (fragments per kilobase of exon per million fragments mapped) (Table 1). Among them, 3556 genes were changed more than or equal to 2-fold with p-values<0.05 in infected cells as compared to non-infected cell. This selection of differentially expressed genes is strict. Very limited changes in RNA expression occurs during the early phases. Only 74 and 223 genes showed significant differential expression at 6 and 12 hpi, respectively. Most expression changes took place at 24 hpi when infection proceeded into the late phase, 2239 and 3060 genes were differentially expressed at 24 and 36 hpi, respectively. Fewer differentially expressed genes were detected in this study as compared to our early study, in which 1267 and 3683 cellular genes were selected as differentially expressed at 12 and 24 hpi. However, the former study was less stringent and included gene covered with more than 1 reads (42).

**Table 1.** In total 12,927 cellular mRNAs were detected in five time points together. Among them 9,738 mRNAs were common between all time points. Expression of 6,860 mRNAs reached to a significant level with a minimum of 10 FPKM. Among them 3556 genes expressions were changed ≥2-fold in infected cells as compared to non-infected cells. Numbers of mRNAs at each time point are listed here.

| **Selection** | **Mock** | **Ad2-6 hpi** | **Ad2-12 hpi** | **Ad2-24 hpi** | **Ad2-36 hpi** |
| --- | --- | --- | --- | --- | --- |
| **$\geq 1$ FPKM** | 11064 | 11163 | 11837 | 11426 | 11191 |
| **$\geq 10$ FPKM** | 5001 | 4846 | 5184 | 4692 | 4371 |
| **$\geq 2$-Fold change** |  | 74<sup>a</sup> | 223 | 2239 | 3060 |
| **Up-regulated** |  | 65<sup>b</sup> | 138 | 1694 | 2142 |
| **Down-regulated** |  | 9<sup>c</sup> | 85 | 545 | 918 |

^a^Number of genes expression with or decrease in Ad2-infected as compared to uninfected cells as measured by sequence reads.

^b^Number of genes expression with more than 2-fold increase in Ad2-infected as compared to uninfected cells.

^c^Number of genes expression with more than 2-fold decrease in Ad2-infected as compared to uninfected cells.

Based on the kinetics of changes of gene expression at different stages of infection, 3451 out of 3556 genes fell into 20 major different expression clusters (Figure 1). The complete list of genes in each cluster is included in supplementary Table S1. At 6 hpi, more than 87% of the differentially expressed genes were up-regulated (cluster 1+2+3+4). Expression of nearly all of these genes reached their highest level at 6 hpi, except two which reached their highest levels at 12 hpi. Then about 80% of them became down-regulated during the late phase of infection (clusters 1 and cluster 2). The rest either remained up-regulated (cluster 4), or were gradually reduced to the basal level in the late phase (cluster 3). Only 9 genes (cluster 5) were down-regulated at 6 hpi and their expression remained suppressed until the late phase.

**Figure 1.**
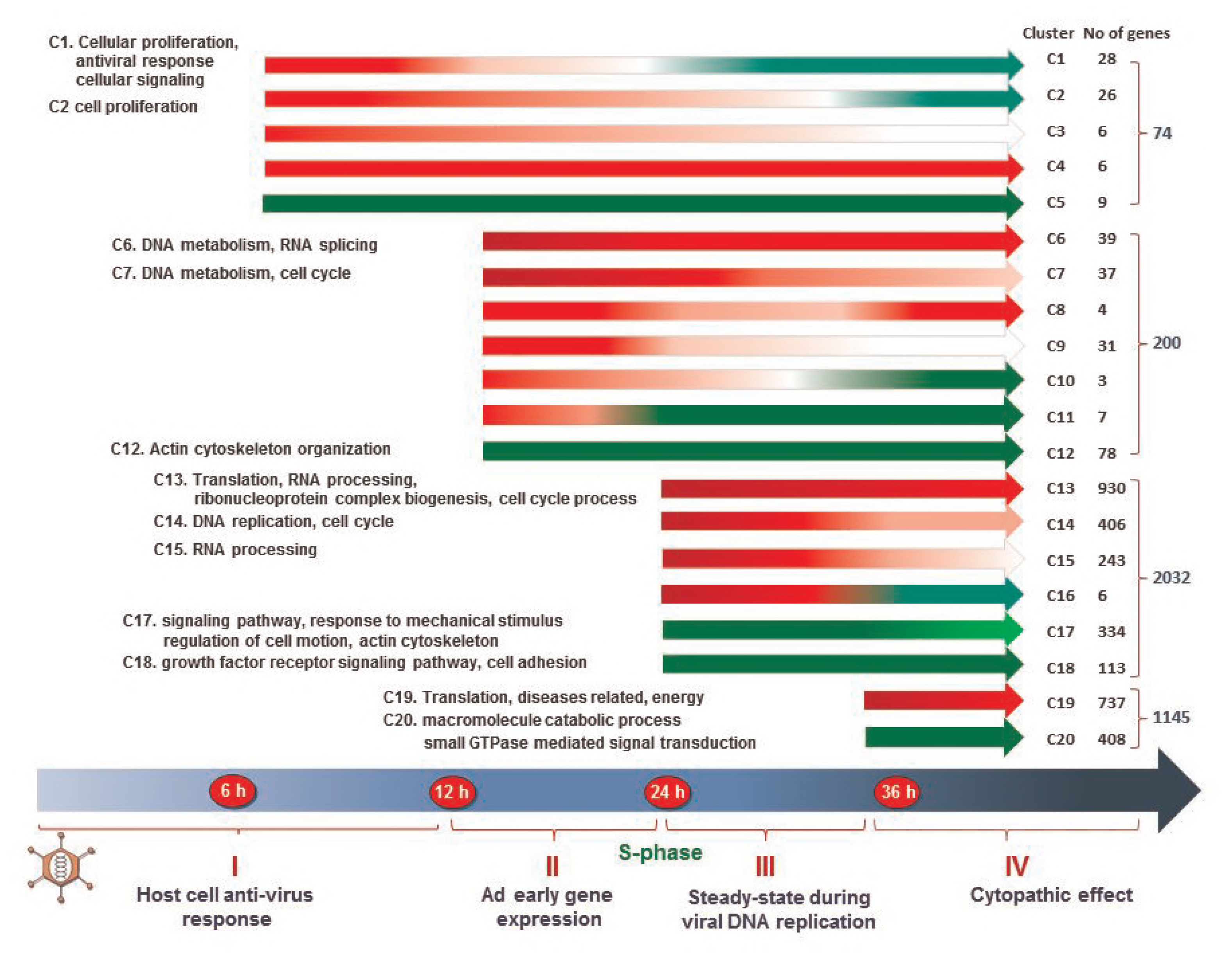
Based on the kinetics of transcription changes, the differentially expressed genes were grouped into 20 clusters (C1-C20). The numbers of genes in each cluster and the number of differentially expressed genes identified at each time point were indicated on the right hand side (Note that the numbers of genes at 12, 24 and 36 hpi are different to the Table 1 because many genes were identified as differentially expressed at more than one time point, but included only in one cluster). The biological functions of genes in each cluster were analyzed using DAVID (left). Red, green and white arrow bars represent RNAs that were up-, down-regulated or unchanged in Ad2-infected cells in comparison to RNA in uninfected control.

At 12 hpi, 122 and 78 genes, became up- and down-regulated in addition to the differentially expressed genes at 6 hpi. Among the up-regulated genes, about 1/3 increased until 36 hpi (Cluster 6), 1/3 remained at a similar level through the rest of the infection (cluster 7), and the remaining 1/3 was only transiently up-regulated at 12 hpi (Cluster 9) and became down-regulated at 24 hpi (cluster 11) or at 36 hpi (cluster 10). Except one gene, all down-regulated genes at 12 hpi remained suppressed until the late phase (cluster 12).

The most dramatic changes in gene expression took place between 12 to 24 hpi then the infection proceeded from the early to the late phase. Thus, expression of 1585 and 447 (2032 in total) additional genes were up- and down-regulated at 24 hpi. Based on the expression changes at 36 hpi, the up-regulated genes at 24 hpi fell into four profiles (Cluster 13+14+15+16). Expression of 59% of these genes increased until 36 hpi (Cluster 13), whereas 25% decreased but remained higher (>2-fold) than in non-infected cells (Cluster 14) and 15% declined to less than 2-fold change at 36 hpi (Cluster 15). Only 6 genes became down-regulated at 36 hpi (Cluster 16). Among 447 down-regulated genes, 75% decreased continually until 36 hpi (Cluster 17), while 25% remained at a similar level (Cluster 18). Change in cellular gene expression was modest between 24 to 36 hpi as compared to that between 12 to 24 hpi. In comparison to non-infected cells, expression of 737 (Cluster 19) and 408 genes (Cluster 20) became up-or down-regulated at 36 hpi in addition to the genes that were differentially expressed at 12 or 24 hpi.

### Biological functions of genes in different expression clusters

The biological consequences of the gene expression changes were analyzed using DAVID (The Database for Annotation, Visualization and Integrated Discovery) and are shown in Figure 1 (left hand panel), and more detailed results are included in supplementary Table S1. The most significant functions of the genes that were transiently up-regulated at 6 hpi (clusters 1-3) were cellular proliferation, antiviral response and cellular signaling. A significant group of genes was cytokines involved stress/immune response and cell growth control. Genes implicated in apoptosis and cell cycle control were also noteworthy. Among transcription factors, up-regulation of ATF3 was the most significant since it reached 6-fold compared to the non-infected control. Expression of ATF3 has been shown to be induced by a variety of signals and is involved in cellular stress response. Only 9 genes were present in cluster 5 and therefore no significant functional categories could be identified by DAVID. However, four (PTPN12, MAP4K3, ERRFI1 and LBH) out of the 9 genes, are involved in cellular signaling and growth control.

During the period between 6 and 12 hpi, adenovirus early genes begin to be expressed, redirecting cellular gene expression. The up-regulated cellular genes are involved in DNA replication (Clusters 6 and 7), including Minichromosome Maintenance Complex Components (MCM) 3, 4, 5, 6, 7 and components of the post-replicative DNA mismatch repair system (MMR) alpha (MSH2-MSH6 heterodimer). In addition, genes implicate in transcription and pre-RNA processing were prominent in cluster 6. Genes implicated in cell cycle were significant in Cluster 7, including CDC25A, CCNE2, CCNE1 and CDK2, the key regulators for the progression from G1 to the S phase. Although no significant function was identified for clusters 8 to 11, several genes, such as JunB, GADD45B and PAPPA function in control of cell growth and proliferation were included in this cluster. The most significant function for the down-regulated genes was actin cytoskeleton organization.

There was a dramatic increase in the number of differentially expressed genes between 12-24 hpi. Cellular genes which function in protein translation became significant among up-regulated genes. These genes covered both cytoplasmic and mitochondrial ribosomal proteins (RPLs/RPSs and MRPLs/MRPSs), eukaryotic translation initiation factors (EIFs), and eukaryotic translation elongation factors (EEFs). Although genes involved in DNA replication and cell cycle were still significant, similar to those at 12 hpi, the number of genes in these categories increased dramatically. For instance, genes involved in DNA metabolism/DNA replication increased from 18 to 124, whereas genes implicated in cell cycle increased from 19 to 153. Most of these genes were included in clusters 13, 14 and 15. The genes involved in DNA replication included DNA polymerases, replication factor C (RFC) and histones from all five families. The large number of genes involved in the cell cycle included many key regulators, such as E2Fs, cyclins, cyclin dependent kinases and cell division cycle (cdc) genes. In addition, genes participating in RNA processing became significant, comprising RNA helicases, the splicing factors, U2 small nuclear RNA auxiliary factors, U3 small nucleolar ribonucleoprotein, U6 small nuclear RNA associated, pre-mRNA processing factors (PRPFs), heterogeneous nuclear ribonucleoproteins (HNRNPs) and small nuclear ribonucleoproteins (SNRNPs). Furthermore, the THO complexes 3, required for efficient export of polyadenylated RNA, were up-regulated, as were the cleavage and polyadenylation specificity factors (CPSFs), playing a key role in the 3’ end cleavage of pre-mRNAs and polyadenylation. Several important components of the exosome complex involved in the degradation and processing of a wide variety of RNA species were also up-regulated.

The number of down-regulated genes between 12 and 24 hpi also increased (clusters 17 and 18). The most significant function of these clusters were cellular signaling, such as TGFBR1, TGFBR2, BMPR2, ACVR1 and SMAD3 involved in TGFβ signaling, as well as growth factors, like EGFR, DGFRA, HBEGF, PDGFC, PLAT, TXNIP, ZFAND5, ARID5B, BCAR1, PDGFRA, PDGFC, VEGFC, NRP1 and PDGFRB. Cytoskeleton organization was significant for genes in cluster 17, whereas genes implicated in cell adhesion were significant in cluster 18.

Previous experiments have shown that the replication of Ad2 DNA reaches a maximum rate during the period from 24 to 36 hpi (36). However, cellular gene expression was still maintained at a high level. The most significant function of the up-regulated genes (cluster 19) was protein translation similar to that at 24 hpi, but with an increased number of genes. Genes involved in the generation of precursor metabolites and energy, as well as oxidation reduction became significant. In addition, several genes identified in different diseases were also significant. The major function for the down-regulated genes (cluster 20) was cellular macromolecule catabolic processes such as ubiquitination and subsequent proteasomal degradation of target proteins. Another significant function was small GTPase mediated signal transduction, involved in vesicle transport.

### Consensus transcription factor binding sites in the promoter region of genes in the different clusters

Genes sharing a similar transcription profile are likely to be regulated by the common transcription factors (TF) or TFs from the same family. To this end, the genes in the 20 different clusters were subjected to analysis for the presence of consensus TF binding sites in their promoter regions (−300 to +100) using Transfind (56). The top ten of the over-represented TF binding sites are listed in the order of significance and are included in the supplementary Table S2. NFκB and c-Rel binding sites were most significant for the genes in cluster 1. Interesting genes among them were BIRC3, IKBA, IL8, CCL20, GROA (CXCL1), TNAP3 and TNF15. These genes are known to be involved in immune response or apoptosis. No significant enrichment of TF binding sites was identified for the genes in clusters 2, 3, 4, 5. For the genes in clusters 6 and 7, only the E2F binding site was significant. Genes with E2F binding sites became more significant in clusters 13, 14 and 19. In addition, the binding sites for GABP, NRF1, and ATF/CREB family were significant among genes in clusters 13, 14 and 15. GABP regulates genes that are involved in cell cycle control, protein synthesis, and cellular metabolism. NRF1 activates the expression of key metabolic genes regulating cellular growth. The ATF/CREB family has diverse functions in controlling cell proliferation and apoptosis. In contrast, the TF binding sites among the down-regulated genes were less significant. Only the MZF1 and AP2 binding sites were scored but their significance was low and they were only present on 8 or 7 genes, respectively. MZF1 can function as a tumor/growth suppressor and controls cell proliferation and tumorigenesis (57). At 36 hpi, different sets of TF binding sites became significant for up-regulated genes (cluster 19), including SP1, STRA13 and NF-Y in addition to GABP while the binding sites for E2F became less significant. This correlated very well with the expression profile of E2Fs. Expression of all E2Fs increased at 12 and 24 hpi, and then decreased at 36 hpi. The TF binding sites for the down-regulated genes were less significant and STRA13 and USF were on the top of the list. STRA13 is a transcriptional repressor. Correspondly, its expression increased 4 and 8 times at 24 and 36 hpi, respectively. STRA13 is involved in DNA damage repair and genome maintenance. Surprisingly, the STRA13 binding site was significant for both up- and down-regulated genes at 36 hpi. Its transcriptional repression is probably mediated by recruitment of other regulatory factors and, depending on the cofactors, STRA13 plays divergent roles. USF that binds to a symmetrical DNA sequence (E-boxes; 5-CACGTG-3) is involved in the transcriptional activation of various genes implicated in physiological processes, such as stress response, immune response, cell cycle control and tumor growth.

### Inconsistency between changes in RNA and protein expression highlighted in genes involved the cellular immune signaling pathway

About 35% of genes that were expressed at the RNA level here were detected at the protein level at 24 and 36 hpi as shown in our previous study using high throughput SILAC-MS technology (Zhao et al 2017). Among 2648 and 2394 proteins that were detected at 24 and 36 hpi, expression of 659 and 645 protein were changed ≥ 1.6-fold as compared to the uninfected control. The correlation between changes in RNA and protein expression was surprisingly low (r=0.3). The functions of the discordantly expressed proteins were analyzed using the web-based tool DAVID, a functional enrichment analysis by intergrating a wide-range heterogeneous data content. However, this tool is less specific for analysis data of virus-induced changes in gene expression because of under representive of genes related to virus infection. In addition, our previous analysis included only proteins that had minimum fold-change of 1.6 and proteins with a slow kinetic of synthesis or degradation might have escaped detection. Nonetheless, we noted that several genes which showed opposite profiles for RNA and protein expression were involved in immune response. Here, we have extended our comparison between RNA and protein expression on genes involved in cellular immune pathways including new SILAC-MS data for 6 and 12 hpi (manuscript submitted for publication). Significantly, many key regulators in cellular immune pathways, NFκB, STAT, apoptosis and MAV displayed inconsistent expression profiles between RNA and protein expression as listed Table 2 and their expression profiles are shown in Figure 2.

**Figure 2.**
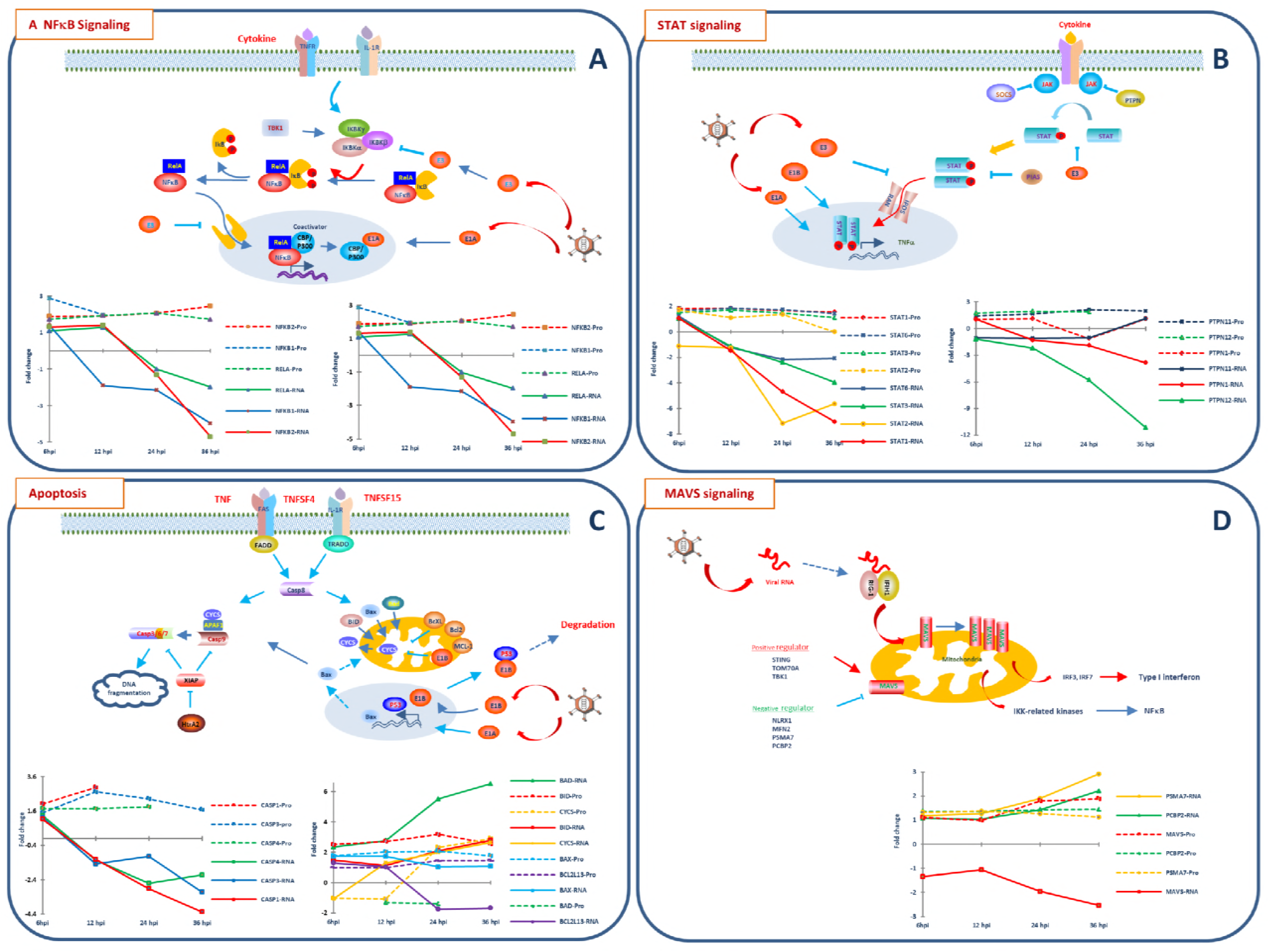
Schematic representation of key components of NFκB (A), STAT (B), apoptosis (C) and MAVS (D) pathways. The involvement of Ad2 proteins is also indicated. Graphs display the expression profiles (fold change between Ad2 and mock) of genes that detected at both protein (dash line) and RNA (solid line) levels at 4 time points. The same color is used for the corresponding protein and RNA.

**Table 2.**
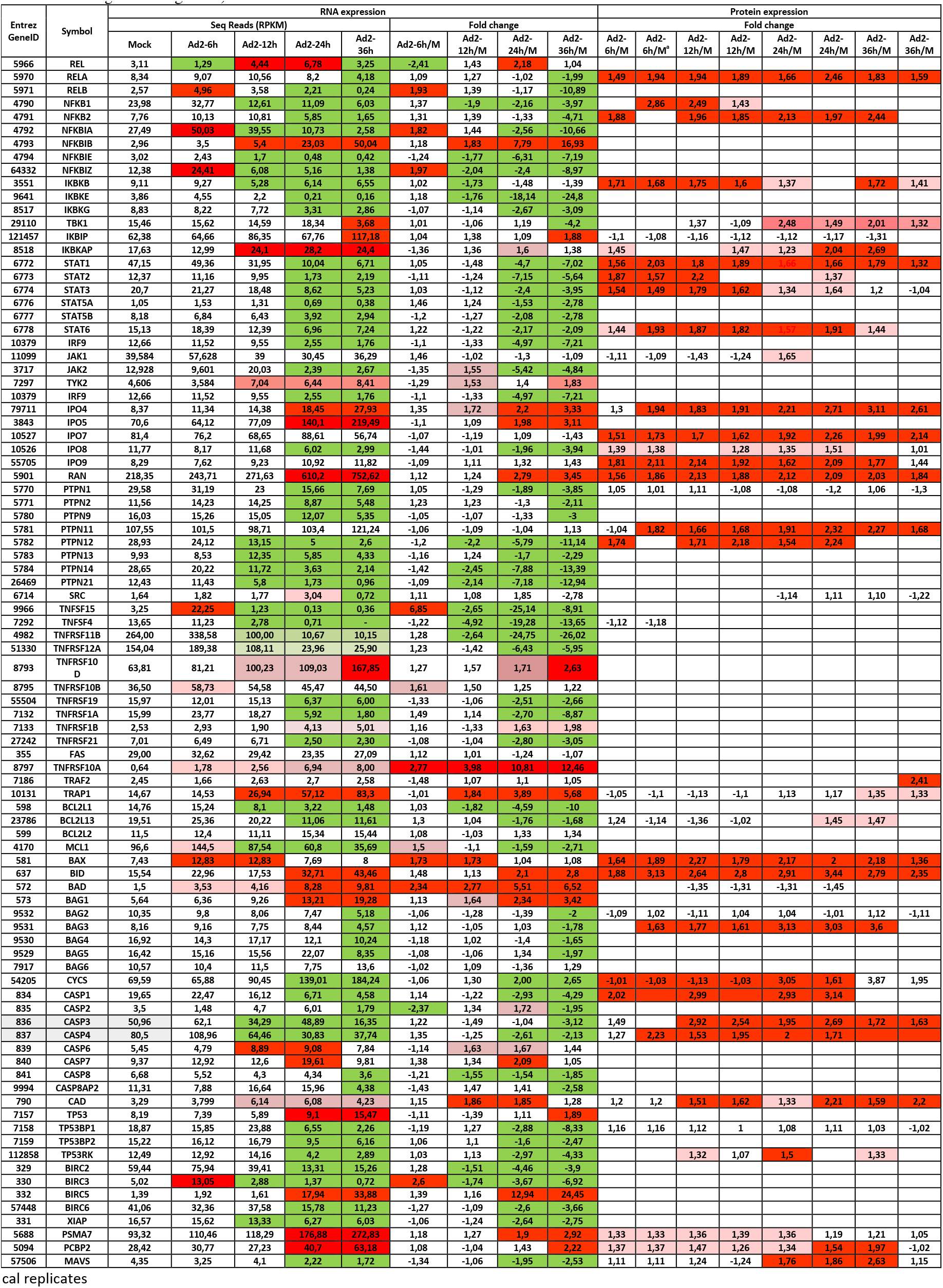
Expressions of genes involved in cellular immune pathways at the RNA and protein levels (Up- and down-regulated RNAs or proteins were highlighted with red or green background).

^a^Biological replicates

As presented above, NFκB and c-Rel binding sites were the most significant in the promoter region of genes that were transiently up-regulated during the early phase. Indeed, expression of several key factors of the NFκB pathway were significantly changed at both the RNA and protein levels (Figure 2A). The transcription of all NFκB family members was detectable, and NFκB1 was the most highly expressed. Except REL, all showed very similar expression profiles. Specifically, they were moderately induced during the early phase, but decreased rapidly and became down-regulated after 24 hpi. Among them, expression of RELA, NFκB1 and NFκB2 was also detectable at the protein level. Coupled with the increased RNA level at 6 hpi, these proteins were all up-regulated. Unexpectedly the NFκB2 and RELA protein levels remained constant until the late phase of infection in spite of the reduction in transcription. The members of NFκB inhibitor family (IκB) displayed diverse transcription profiles. NFKBIA (IκBα) and NFKBIZ (IkBζ) were the most highly expressed and showed similar expression profiles, transiently up-regulated at 6 hpi but decreased at12 hpi and were reduced more than 8-fold at 36 hpi. NFκBIB (IκBβ) showed an opposite expression pattern, low in uninfected cells and at 6 hpi, but increased after 12 hpi and became up-regulated more than 16-fold at 36 hpi. Thus, it appears that NFKBIB replaced NFKBIA to be the most highly expressed IκB in the late phase. None of these gene products was detected at the protein level. The expression changes of the inhibitors of NFκB kinases (IKKs) subunit, IKBKB, and its regulatory subunit IKBKG, as well as IKK-related kinases, IKBKE and TBK1, appeared to be coordinated. They were delayed as compared to the expression of NFκBs and IκBs and significant down-regulation of transcription occurred at 24 or 36 hpi. A surprising finding was that the expression of IKBKB and TBK1 was up-regulated at the proteins level. The IKBKB protein was up-regulated already at 6 hpi and remained stable until the late phase while the up-regulation of the TBK1 protein was significant after 24 hpi. The results thus indicate that the positive regulators of the NFκB pathway are activated at both the RNA and protein levels during the early phase of infection as result of the host immediate response to the infection. Following the progression of the infection, these proteins remained up-regulated until 36 hpi although their transcription was suppressed. However, the fact that the downstream target genes of the NFκB pathway were down-regulated during the late phase indicates that these proteins have lost their functions as transcriptional activitors. The dramatic up-regulation of NFKIB may contribute to the inhibition of the NFκB activity. Other post-translational control mechanisms, such as the blocking of the nuclear transport, loss of its coactivator such as CBP/P300, p400 and TRAPP due to interaction with the Ad2 E1A protein, may contribute to the block of the NFκB activity (58, 59). Other yet unidentified mechanism might also cause the inactivation of the NFκF pathway.

The Janus kinase-signal transducer and activator of transcription (JAK/STAT) signaling is another important pathway regulating the innate immune response. Transcription of all STATs (STAT1, STAT2, STAT3, STAT5A/B and STAT6) was unchanged up to 12 hpi, but were then down-regulated after 24 hpi. Four STAT proteins (STAT1, STAT2, STAT3 and STAT6) were detected and they were up-regulated during the early phase and remained stable or decreased slightly in the late phase. JAKs are important activators of STAT and catalyse the phosphorylation of the STAT proteins. The three JAK kinases, JAK1, JAK2 and TYK2, displayed different expression profiles. JAK1 was the most highly transcribed and only slightly increased at 6 hpi. Then, it decreased to the basic level and remained constant until the late phase. Transcription of both JAK2 and TYK2 increased at 12 hpi. JAK2 decreased during the late phase while TYK2 remained constant. Only JAK1 protein was detected and it decreased slightly during the early phase, but became up-regulated at 24 hpi. The activity of the STAT proteins is also controlled by several negative regulators, including protein tyrosine phosphatase (PTPN), suppressor of cytokine signaling (SOCS) and protein inhibitor of activated STAT (PIAS). Several PTPNs were detected at both RNA and protein levels with inconsistent expression profiles. Either their RNAs were down-regulated, while their protein levels remained constant (PTPN1) or increased (PTPN12). In other cases the RNA remained stable, while the protein level was increased (PTPN11). Furthermore, several Importins and Ran, required for nuclear translocation of STATs, were up-regulated at both at the RNA and protein levels during the infection. Suppression of STAT transcription and promotion of STAT translation during Ad2 infection might reflect an aspect of the battle between the virus and its host. Through several distinct routes the infected cells recognize different viral components and active the expression of IFNβ which leads the stimulation of the JAK/STAT pathway. In turn, viruses have developed strategies to circumvent the IFNβ response (60). Expression of most, if not all, of the downstream targets of the STAT pathway were suppressed, indicating that the STAT pathway is blocked (data not shown). Apparently, the inhibition of the STAT pathway occurs both transcriptional and post-translationally. The activity of STATs has been shown to be modulated by various posttranslational modifications (61, 62). Upon infection, adenovirus uses several strategies to block the STAT pathway. The viral E1A plays a significant role in the inactivation of the STAT pathway by binding to STATs, or their coactivator CBP/p300.(11, 63–65). Meanwhile, the E1B-55k protein represses expression of IFN-inducible genes which leads to the inhibition of the STAT signaling pathway (66). In addition, E3-14.7K protein interacts with STAT1 which results in the inhibition of STAT1 phosphorylation and nuclear translocation (67). Furthermore, phosphorylated STAT1 has been shown to be sequestered at viral replication centers in the nuclues (68).

Apoptosis pathways are extensively regulated during Ad2 infection. Our RNA sequencing results showed that transcription of more than 60% of genes that are directly involved in apoptosis were down-regulated, whereas only 20% were up-regulated in the late phase (data not shown). Transcription of most TNF family ligands was undetectable or at a very low level except for TNFSF15 and TNFSF4 (Table 2). Both of them decreased after 12 hpi, although TNFSF15 was transiently induced more than 6-fold at 6 hpi. Numerous TNF receptor superfamily members were expressed at the transcriptional level with diverse expression profiles. TNFRSF11B and TNFRSF12A were the most highly expressed receptors. After being slightly increased at 6 hpi, their RNA levels decreased at 12 hpi and were then more than 25- and 6-fold lower at 36 hpi as compared to the non-infected control. Unfortunately, none of the TNF receptor superfamily members was detected at the protein level.

Caspases (CASPs) and the Bcl2 families are key player in apoptosis. At the transcriptional level, CASPs showed different expression profiles. Among them, CASP1, 3 and 4 are most highly expressed with similar expression profiles, slightly increased at 6 hpi and then down-regulated. All of these CASPs were detected at the protein level and were up-regulated, opposite to their RNA expression profile. Expressions of most Bcl2 family members were low at the RNA level except for those listed in the Table 2. Transcriptions of most anti-apoptotic BCLs (BCl2A1, BCL2L1, BCL2L13 and MCL1) were down-regulated after a slight increase at 6 hpi. Among them, only BCL2L13 protein was detected which showed 40% increased expression during the late phase. Among pro-apoptotic genes, transcription of BID, BAD and BAX was up-regulated gradually towards the late phase or remained stable. At the protein level, BID and BAX were up-regulated from the early to the late phase, although the RNA level for BAX decreased in the late phase. BAD protein displayed an expression pattern opposite to its RNA.

Although most genes that are directly involved in apoptosis were down-regulated at the transcriptional level, several important pro-apoptotic players were up-regulated at the protein level (CASP3, BAX and BID). The fact that apoptosis is efficiently inhibited during an adenovirus infection indicated that the functions of these proteins must be inactivated. To counteract the host defensive apoptotic pathways, adenoviruses have established very efficient mechanisms by encoding their own anti-apoptotic proteins in the E1B and E3 regions. Obviously, the regulation of apoptosis is very multifaceted and a comprehensive view of how apoptosis is blocked during an Ad2 infection remains to be unraveled.

Inconsistent expression profiles between RNA and protein for the genes involved in MAVS was shown in our previous study (39). We show here that expression of MAVS is stable at both RNA and protein levels during the early phase, whereas difference were seen in the late phase. In addition, we have studied the expression of three MAVS regulatory proteins, PSMA7, PCBP2 and TBK1. The expression profiles of the negative regulators PSMA7 and PCBP2 were similar at both the RNA and protein levels, and increased slowly during infection. The positive regulator, TBK1 showed an opposite profile; its RNA was down-regulated at 36 hpi whereas its protein level increased after 24 hpi. In spite of the up-regulation of MAVS and its positive regulator, expression of its target genes (type I interferon genes) was very low or undetectable, suggesting that this antivirus pathway is inactivated during the late phase.

Furthermore, different expression profiles were also observed for galectins LGALS. LGALS3 and 8 (Gal3 and 8) were the most highly expressed among LGALSs and their RNAs were down-regulated after 24 hpi. However, their proteins remained constant from early to late phase. Galectins have been shown to involve in innate immune processes (69). Colocalization of LGALS3 with incoming Ad5 has been observed and its role in Ad5 transport was suggested (70). Stable expression of LGALS3 has been reported previously in Ad5-infected cells, while it is down-regulated in Ad3-infected cells (71).

Apparently, there are features common to genes which participate in important immune pathways, being suppressed at the transcriptional level while being enhanced or stable at the protein level during the late phase. Thus, we hypothesize that the transcriptional regulation of these genes is mediated mainly by Ad2, while protein translation machinery is not yet completely under the control of the virus. The increased protein expression could be caused by increased translation and/or decreased protein degradation. The facts that most of downstream genes of immune pathways are down-regulated at RNA level in the late phase even though their key regulators are stable or up-regulated at the protein level. Apparently, adenovirus-mediated post-translational mechanisms play a very important role. As discussed above, inhibition of STAT pathway represent the best example how adenovirus has evolved redundant strategies to counteract cellular immune response. By regulating protein modification, blocking of protein-protein interactions, inhibiting of the protein transport to its destination, or direct interacting, adenovirus controls host cell antiviral pathways. Last but not least, non-coding RNAs (ncRNAs) have been shown to be important regulators of various biological processes. Alternations of cellular miRNA and lncRNA expression during Ad2 infection have been studied using RNA-seq (37, 38). Significant changes in their expression take place after 24 hpi. The strong correlation of ncRNA expression changes with infection progression indicates that ncRNA play important roles during infection. Specially, majority of differentially expressed miRNAs were down-regulated in the late phase. One major mechanisms of miRNA in gene regulation is the suppression of translation through partial complementary to 3’ UTRs of mRNA. Thus, down-regulation of miRNAs could lead to stable or increase translation of special sets of proteins. In contrast, most of differencially expressed cellular lncRNAs were up-regulated in the late phase. Several lncRNAs that are predicted to target immune response genes were down-regulated in the late phase. In addition, a large share of differentially expressed lncRNA are associated with RNA-binding proteins (RBPs), being involved in posttranscriptional RNA processing and translation regulation. However, how they are regulated, and how they are involved in the regulation of cellular gene expression during adenovirus infection nees to be further addessed.

## Acknowledgments

Sequencing was performed at the SNP&SEQ Technology Platform at Uppsala University and University Hospital. We thank Ulrika Liljedahl and Johanna Lagensjö for excellent sequencing. Martin Dahlö at UPPMAX is acknowledged for helping to make the code run on the UPPMAX resources. We thank Dr. Caroline Gallant for critical reading of the manuscript. This work was supported by the Kjell and Märta Beijer Foundation (UP), Åke Wiberg Foundation (SBL) and Magnus Bergvall Foundation (SBL).

